# Fiber optical parametric amplification of low-photon-flux microscopy signals

**DOI:** 10.64898/2026.03.25.714345

**Authors:** J. Demas, L. Tan, S. Ramachandran

## Abstract

The performance of a laser scanning microscope inevitably depends on the performance of the point detector. As laser scanning approaches aim to penetrate deeper in tissue, there is a commensurate need for detectors that can operate with high sensitivity, bandwidth, and dynamic range at near-infrared wavelengths where scattering is reduced. Here, we demonstrate that fiber optical parametric amplification can be used to boost low-power microscopy signals to levels that can be detected by near-infrared photodiodes without introducing prohibitive noise. We construct amplifiers that achieve >50 dB of parametric gain at wavelengths within the third near-infrared transparency window and have similar sensitivity to near-infrared photomultiplier tubes. Furthermore, these amplifiers outperform detection with a photodiode and subsequent electrical amplification, providing a factor of 10–100-fold improvement in sensitivity. We demonstrate amplifier bandwidths up to ~1.6 GHz, a factor of 10 faster than conventional detectors, including near-infrared photo-multiplier tubes, with sensitivity of ~8 nW (corresponding to ~20 photons/pixel). Finally, the increased performance of the optical amplifier is confirmed in diagnostic imaging experiments where >10× less power is required to achieve the same signal-to-noise ratio and contrast as images using electrical amplification. Accordingly, fiber optical parametric amplification is a new path forward for extending the performance of laser scanning microscopes in the near infrared.

## 1. Introduction

Laser scanning microscopy (LSM), in its myriad forms (including optical coherence tomography (OCT) [1], confocal fluorescence microscopy [2], and multiphoton microscopy (MPM) [3], among others) has become essential for characterizing biological tissues [4]. LSM is characterized by point-by-point acquisition of pixels to form images, in contrast to parallel acquisition with a camera or detector array – as in conventional widefield microscopy. The information comprising each pixel is measured serially by a single point detector, limiting imaging speeds to slower frame rates than parallel approaches. However, LSM retains unique advantages, such as optical sectioning and enhanced resolution, that drive its utility.

LSM has also gained traction as many of its associated techniques can image deeper into scattering tissue than other optical imaging methods. Increased penetration for LSM is facilitated by compatibility with longer wavelength excitation and/or detection because the scattering length in opaque tissue increases with wavelength [5]. Accordingly, the second and third near-infrared (NIR) tissue transparency windows (where scattering is reduced relative to visible light and the absorption spectrum has several local minima) have become loci for deep tissue optical imaging methods including OCT (*λ* ~ 1.3 μm) [1] and three-photon microscopy (3pM, *λ* ~ 1.3 and 1.7 μm for green and red fluorophores, respectively) [6]. Additionally, there are many active efforts to develop NIR fluorescent labels using both organic molecules and bio-compatible quantum dots to facilitate deep tissue confocal fluorescence imaging [7]. Finally, in the last decade, mid-infrared and shortwave infrared photothermal microscopy (MIP and SWIP, respectively) have emerged as techniques for conducting chemical-specific imaging deep in scattering media [8,9]; both techniques use NIR probe beams for deeper penetration.

In any LSM modality, the specifications of the point detector (sensitivity, bandwidth, *etc*.) are crucial to the system performance. Perhaps the most used detectors for LSM are photomultiplier tubes (PMTs) as they are sensitive to single photons, have analog output current proportional to the incident photon flux, and have bandwidths in the hundreds of MHz regime [10]. However, their sensitivity drastically reduces for NIR wavelengths due to low quantum efficiency, and they typically have low, damage-limited dynamic range. The dynamic range in turn limits the effective bandwidth: for a conventional PMT (such as the popular Hamamatsu H11706-40), assuming on average ~2 photons/pixel – the minimum criteria to achieve a signal-to-noise ratio (SNR) above unity – the pixel rate must be below ~10 MHz to stay within the manufacturer-specified output current limit [11]. Thus, the dynamic range constricts the effective bandwidth by >10-fold relative to the device response time.

Single photon avalanche photodiodes (SPADs) are an alternative for point detection, with the advantages being that they have single photon sensitivity and operate at NIR wavelengths [12]. However, these detectors are typically digital – as opposed to analog – in that they can only detect the presence or lack of a single photon rather than resolve the number of incident photons. While some photon-number-resolving SPADs exist, typically they have dynamic ranges of only 3-5 photons [13]. As a result, many repeated measurements are necessary to build reasonable image SNR, placing an upper bound on frame rate. Silicon photomultipliers (SiPMs) are essentially arrays of SPADs that act in parallel as a single point detector, allowing these detectors to produce effectively analog signals by aggregating digital responses, albeit at the cost of increased dark noise relative to PMTs or SPADs [14]. Currently, SiPMs have bandwidths approaching that of PMTs, but have only been commercially realized on silicon, limiting operation to wavelengths of ~1.1 μm and below.

Finally, conventional photodiodes (PDs) can provide exceptional bandwidth, dynamic range, and noise characteristics over a broad range of NIR wavelengths, making them the default choice for NIR detection in many methods, including OCT, MIP, and SWIP [1,8,9]. However, typical PD responsivity is relatively low (~1 A/W), and thus for small signal applications, readout noise becomes prohibitive. PDs (and PMTs, for that matter) are often paired with electrical amplifiers to boost detected signals above the noise floor of readout electronics. However, *post facto* electrical amplification typically limits both bandwidth and SNR. Thus, PDs are ideal detectors for LSM applications with relatively high signal power but low responsivity inhibits their use in low-photon-flux applications.

Here, we attempt to overcome this responsivity gap by pre-amplifying NIR signals from a laser scanning microscope. Inspired by methods commonly employed in telecommunications, we use distributed amplification in optical fibers to increase signal photon flux such that detection with conventional PDs becomes tenable. Parametric amplification maintains the spatial and temporal phase coherence of the signal light and can operate with low SNR degradation, limited only by fundamental quantum noise, and is thus compatible with imaging small signals in LSM applications that require coherence (such as confocal microscopy or OCT) [15]. We demonstrate a sensitivity of ~200 fW/Hz^1/2^, corresponding to a noise equivalent power (NEP) of ~2 nW at a bandwidth of ~100 MHz for a 1.6 μm signal photon wavelength, meeting the theoretical performance of a NIR PMT and exceeding that of conventional electronic amplification by >10-100×. Additionally, we show a sensitivity of ~8 nW at ~1.6 GHz bandwidth, >10× faster than typical electrical amplifiers or NIR PMTs. These advantages in performance relative to electrical amplification are also present in confocal imaging experiments where contrast and SNR are improved by >10-fold. Furthermore, these amplifiers are fiber-based and thus drop-in compatible for alignment-free operation with any existing fiber-based LSM. While this preliminary demonstration does not yet have shot-noise-limited performance, it represents a new paradigm for detection of NIR signals in LSM.

## 2. Parametric amplification in highly nonlinear optical fiber

### 2.1 Parametric amplification

Optical parametric amplification describes a nonlinear interaction between light waves that transfers energy to new frequencies [16]. Parametric processes do not rely on transitions between electronic or vibrational states in a medium and thus are limited by upper state lifetimes; instead, they employ “virtual states” such that interactions can occur instantaneously, allowing for generation of pulses as short as attoseconds [17]. Consequently, parametric processes are ideal for high bandwidth amplification. Furthermore, because parametric gain depends on the optical properties rather than solid state properties of a medium, it is commonly used to amplify wavelengths outside the bandwidth of common solid state laser materials. Optical parametric amplifiers (OPAs) based on bulk nonlinear crystals can generate tunable lasers at exotic wavelengths across the visible, NIR, and infrared spectral regions for myriad applications including MPM, SWIP, Raman spectroscopy, and coherent anti-Stokes Raman spectroscopy (CARS). Parametric nonlinearities also exist in amorphous materials, such as silica glass, which can be formed into waveguides and optical fibers. Accordingly, fiber optical parametric amplifiers (FOPAs) make use of guided modes to increase interaction lengths for higher conversion efficiency. FOPAs have been used for distributed amplification [15] and multi-casting [18] amongst other telecommunications applications [19], as well as to generate high power lasers at novel colors [20,21] and to generate entangled photon pair sources for quantum optical applications [22,23].

Additionally, parametric amplification has the potential to operate with very low noise. Like other optical amplifiers, such as erbium doped fiber amplifiers (EDFAs), parametric amplifiers can operate with noise factors approaching the fundamental quantum limit of *F* = 2 [24]. Here, *F* describes the degradation in SNR due to the amplification process and is fundamentally limited by the equal probability of an excited state (or virtual state) decaying via spontaneous emission (creating a noise photon) or stimulated emission (creating a signal photon).

Low noise optical parametric amplification for imaging has previously been explored using bulk nonlinear crystals operating at NIR wavelengths [25,26]. These demonstrations showcase exceptionally high gain (90 dB, or 10^9^) and sensitivity (~23 pW at 5 MHz bandwidth). However, OPA in crystals requires tight focusing for high intensity, which limits interaction lengths due to the diffraction of the light beams. As a result, both high peak power for efficiency over short lengths and low average power to avoid bulk dielectric damage are required — a regime best satisfied by using ultrafast pulses. Indeed, OPA-based imaging amplification has only been demonstrated in the ultrafast regime thus far. The need for multiple ultrafast pulse trains at different wavelengths necessitates a complicated and unwieldy experimental apparatus where inter-pulse timing and dispersion management become critical. Furthermore, parametric gain only occurs while pulses overlap in time. Therefore, this scheme would be incompatible with amplifying fluorescence signals with finite lifetimes (~ ns) or time-of-flight signals such as those generated by OCT (~ ps). Finally, the bandwidth is limited by the repetition rate of the ultrafast laser, typically ~100 MHz at fastest; thus, throttling speeds to >10-fold slower than the capabilities of a conventional NIR PD.

In contrast, a FOPA-based image amplifier can have a long interaction length due to the guided wave geometry of optical fibers, allowing for amplification with long duration pulses and even continuous wave (CW) beams. CW operation allows for a simpler setup, is compatible with amplifying long duration signals, and can use the entire bandwidth provided by the PD. Finally, FOPAs can be configured in a spliced, all-fiber format that is drop-in compatible with fiber-based LSMs and does not require free-space alignment.

### 2.2 Fiber optical parametric amplifiers

In optical fibers, the relevant nonlinearity is third order, such that four waves interact to facilitate parametric gain – a process often referred to as “four-wave mixing” (FWM) [27]. In FWM, two high power waves give rise to a lower frequency “signal” wave and a higher frequency “idler” wave. Often, the high-power waves are degenerate in wavelength (as is the case here) and treated as a single “pump” beam. Efficient transfer of energy through FWM requires the conservation of energy such that the energy of two pump photons is equivalent to the sum of the energies of the output signal and idler photons. Additionally, momentum must be conserved, requiring “phase matching” between the waves. This phase matching relation, *Δβ*, is given by:

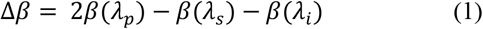

where *β*(*λ*) is the propagation constant of the fiber mode at wavelength *λ*, given by *β*(*λ*) = 2*πn*_*eff*_(*λ*)/ *λ, n*_*eff*_ is the effective refractive index, and the subscripts ‘*p*’, ‘*s*’, and ‘*i*’ correspond to pump, signal, and idler, respectively. Interestingly, accumulation of extra phase at the pump wavelength due to an effect called self-phase modulation [27] offsets Δ*β* such that FWM gain occurs for the condition where *–4γP ≤Δβ≤0* with maximum gain at Δ*β = –2γP*. The factor *2γP* accounts for the pump self-phase modulation, where *γ* is the nonlinear parameter given by *γ = 2πn*_*2*_*/*λ*A*_*eff*_, *n*_*2*_ is the nonlinear refractive index of the medium (*n*_*2*_ ~ 2.7·10^−20^ m^2^/W for silica glass), *A*_*eff*_ is the effective mode area, and *P* is the pump power.

If phase matching and energy conservation can be satisfied simultaneously at a given combination of pump, signal, and idler waves, high power pump light will spontaneously generate signal and idler photons within the fiber – akin to spontaneous emission in a conventional laser. If a low power seed laser tuned to the signal wavelength co-propagates with the high-power pump beam, the signal wave will extract energy from the pump and be amplified. Due to conservation of energy and momentum, a wave at the idler wavelength is necessarily generated and grows alongside the amplified signal in the fiber. At wavelengths where the phase matching and energy conservation conditions are exactly met, the peak of the small signal gain spectrum centered on *λ*_*s*_ is given by:

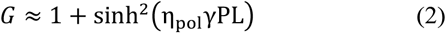

where *L* is the length of the fiber, and *η*_*pol*_ is a factor that accounts for depolarization between the pump and signal over the length of the fiber (*η*_*pol*_ ~ 0.66) [27]. Figure 1(a) shows the calculated phase matching (upper plot) and parametric gain (lower plot) for a 480 m length of highly nonlinear fiber (HNLF). For *λ*_*p*_ = 1546 nm, broadband gain is predicted for a signal band at 1600*–*1650 nm and an idler band at 1450*–*1500 nm. For ~2.5 W of pump power, the simulations predict >50 dB (>10^5^) of parametric gain – on the order of typical PMT gain. Note that because FWM is a coherent process, the phase of the signal is preserved during amplification, and its conjugate phase is imprinted onto the idler wave. Furthermore, because FOPAs are usually constructed with single-mode fibers (as is the case here), only a single spatial mode is guided – thus the parametrically amplified signal and idler waves are both temporally and spatially coherent with respect to the input signal.

**Fig. 1.**
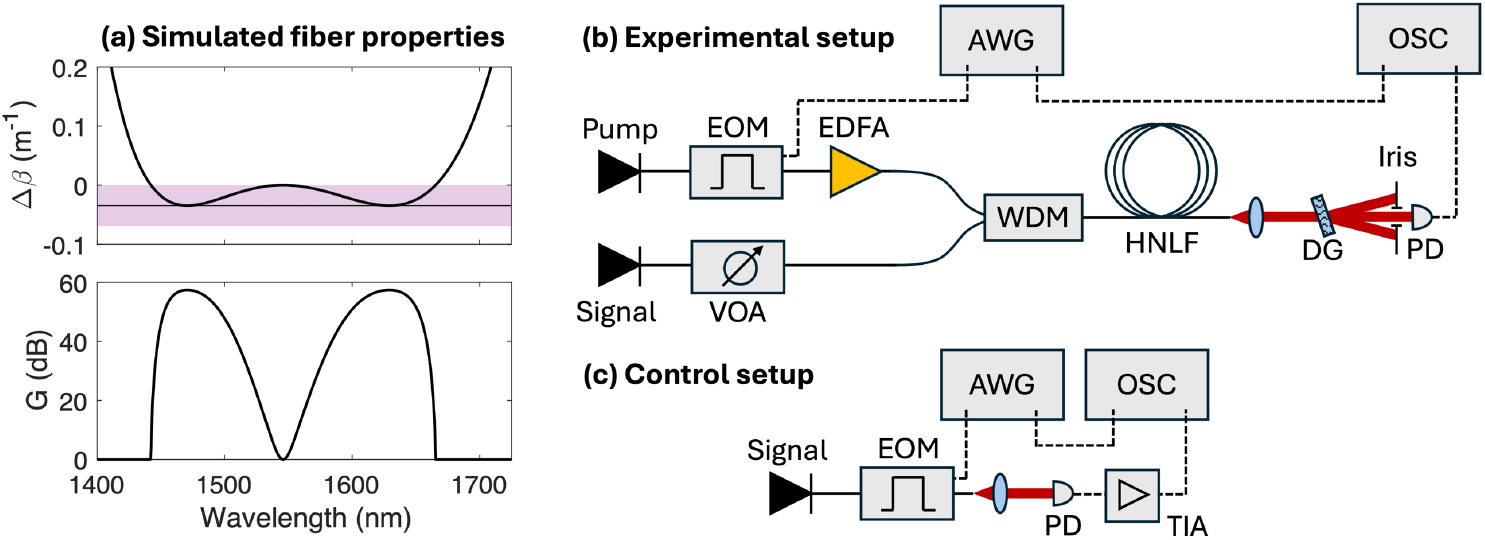
(a) Simulated phase matching and optical parametric gain for HNLF; shaded purple region in the upper plot indicates FWM gain bandwidth corresponding to the peaks in the lower plot. (b) Experimental setup for characterizing optical amplification and detection sensitivity; AWG = arbitrary waveform generator, EOM = electro-optic modulator, VOA = variable optical attenuator, EDFA = erbium-doped fiber amplifier, WDM = wavelength division multiplexer, HNLF = highly nonlinear fiber, DG = diffraction grating, PD = photodiode, OSC = oscilloscope. (c) Setup for characterizing direct detection and electrical amplification; TIA = transimpedance amplifier.

### 2.3 Experimental gain characterization

Figure 1(b) shows the experimental setup for probing parametric gain in HNLF. The pump laser consists of an external cavity diode laser (ECL) tuned to *λ*_*p*_ = 1546 nm and modulated by an electro-optic modulator (EOM) to generate pulses. An arbitrary waveform generator (AWG) drives the EOM with pulse durations varying from Δ*τ* = 0.5*–*16 ns. Pulsing reduces the duty cycle of the pump to reduce the average power requirements for the characterization experiments and to emulate signals with different bandwidths; in a fully realized system, a CW pump would be preferable. Subsequently, an EDFA amplifies the peak power of the pulses to a maximum power of *P* = 25 W.

The signal beam comprises a second ECL (*λ*_*s*_ = 1620 nm) and a variable optical attenuator (VOA) to seed the amplifier with CW laser light over a range of *P*_*signal*_ = 10^*–*2^*–*10^3^ nW. The signal and pump are combined with a wavelength division multiplexer (WDM) which is spliced to a length of HNLF (OFS standard HNLF). We used two different fiber lengths in experiments: 50 m and 480 m. In FWM, the signal and idler waves have the same number of generated photons; thus, both waves are viable for detection. Here we have elected to detect the idler because the blue-shifted wavelength is subject to less potential SNR reduction from parasitic nonlinear processes, such as spontaneous Raman scattering [27], however either (or both) wavelength could be detected depending on the needs of the application.

The fiber output is collimated with an aspheric lens (Thorlabs C110TMD-C), and a diffraction grating (DG) is used to separate the pump, signal, and idler beams. The idler beam is isolated with an iris and detected using an InGaAs PD (Thorlabs DET08C, *Δv* = 5 GHz). The DG-based spectral filter has a passband of ~0.7 nm centered at *λ*_*i*_ = 1479 nm, a loss of 2.6 dB (55% transmission), and serves to remove amplified spontaneous emission (ASE) resulting from the parametric gain process which would otherwise degrade SNR. We found that employing the DG-based filter, as opposed to a conventional dielectric bandpass filter (Thorlabs FBH1480-12, 12 nm passband centered on 1480 nm, >90% transmission), improved the SNR by a factor of ~3, despite causing higher signal loss.

To benchmark the FOPA performance, we also constructed a control setup (Fig. 1c) by directly sending the signal light (*λ*_*s*_ = 1620 nm) through the EOM and onto the PD. After the PD, we installed one of two different (Femto DHPCA-100, Thorlabs TIA60) transimpedance amplifiers (TIAs) during experiments to characterize NEP with electrical, rather than optical, amplification. Signals from both the experimental (Fig. 1b) and control (Fig. 1c) cases were characterized with an oscilloscope (Agilent 86100A, 3.2 GS/s sampling rate).

Figure 2a shows the output spectrum of the FOPA measured with an optical spectrum analyzer (Ando AQ6317). The initial signal light (black line) is amplified in the presence of the pump, and an idler peak is generated on the shorter-wavelength side within the detection band (gray shaded region) defined by the DG filter. ASE from the FWM process manifests as broadband peaks below the signal and idler lines and is the primary source of noise for the FOPA under high gain conditions.

**Fig. 2.**
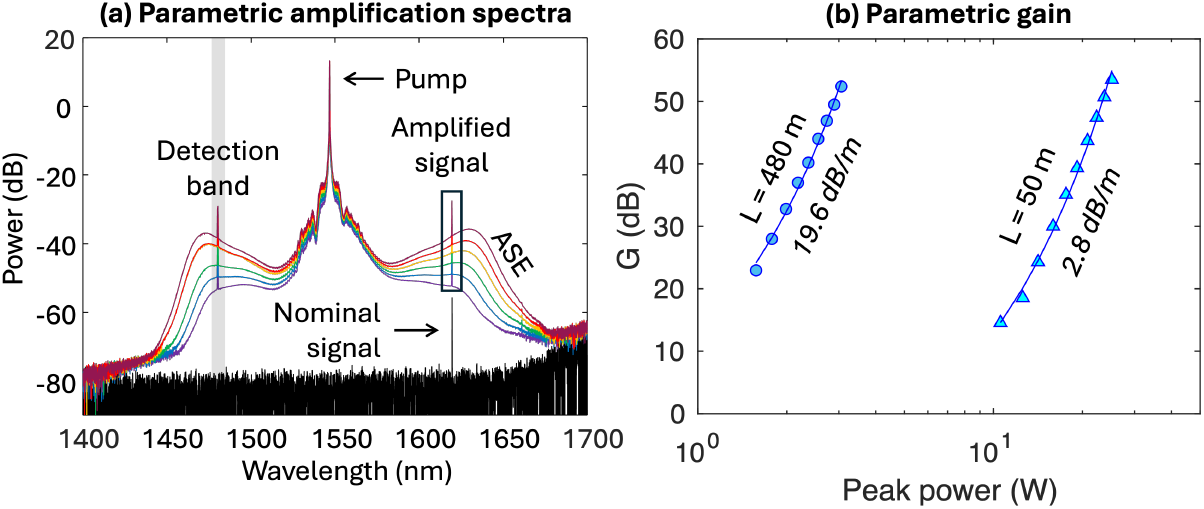
(a) Example optical spectra showing the parametric amplification process; high power pump (*λ* = 1546 nm) is combined with low power signal (*λ* = 1620 nm, black line) in L = 50 m of HNLF; increasing pump power amplifies the signal and creates idler photons (*λ* = 1479 nm) within the PD detection band; amplified spontaneous emission (ASE) creates noise photons outside the signal and idler lines making bandpass filtering necessary to preserve SNR. (b) Signal gain as a function of pump peak power for 480 and 50 m HNLF lengths; longer fiber length requires lower power to achieve parametric gain, while longer fibers provide broader spectral gain bands.

We also characterized the system small signal gain, *G*, by measuring the increase in the average power of the signal and accounting for the duty cycle of the pump pulses (Fig. 2b) for both 50 and 480 m lengths of HNLF. The 50-m amplifier features broad gain peaks (~33 nm full-width at half-maximum, Fig. 2a) but requires higher pump power (~20 W) to produce the desired gain. The 480-m amplifier has narrower gain (~6 nm) and requires less pump power (~3 W), making it more practical for some cases. However, both amplifiers can achieve >50 dB gain; on par with the performance of typical PMTs.

## 3. Measurement of noise equivalent power

The FOPA (Fig. 1b) generates pulses at the idler wavelength due to the parametric interaction between the pump laser and low power signal light. Figure 3a shows example pulses for a 2 ns pump pulse co-propagating with signal powers ranging from 0 to 200 nW. We mimic the function of a field-programmable gate array (FPGA) for signal digitization in a laser scanning microscope by integrating over the duration *Δτ*, defined by the *1/e*^*2*^ full width of the generated idler pulse (purple shaded region, Fig. 3a):

**Fig. 3.**
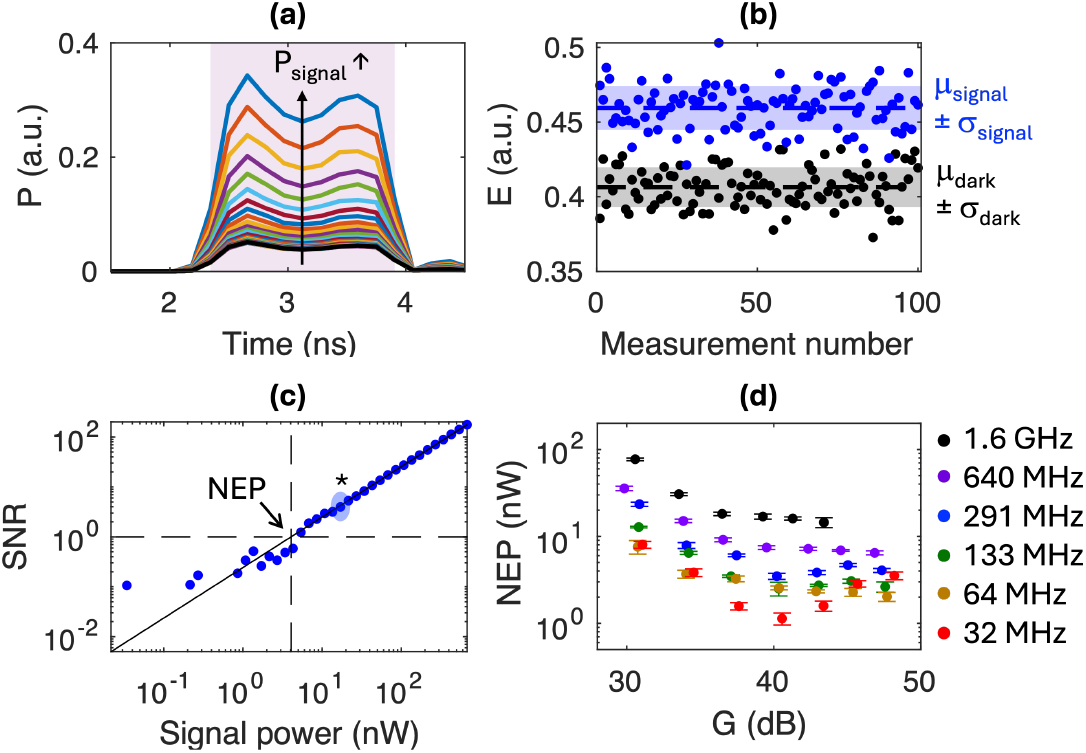
(a) Example idler pulses (*Δτ* = 1.7 ns, Δ*v* = 291 MHz) as a function of increasing signal power. (b) Example measurements of idler pulse energy for *P*_*signal*_ = 17 nW; blue shaded region corresponds to mean (*μ*_*signal*_) ± standard deviation (*σ*_*signal*_) of measurements; gray shaded region corresponds to measurements with no signal present corresponding to amplifier noise (*μ*_*dark*_ ± *σ*_*dark*_); calculated SNR = 4.1. (c) SNR as a function of signal power for 50-m amplifier; measurement in (b) highlighted with blue shading and asterisk; intercept of the fitting line with SNR = 1 corresponds to the noise equivalent power (NEP). (d) NEP as a function of bandwidth and parametric gain; Error bars correspond to 95% confidence interval of SNR fitting line, *i.e*. black line in (c).

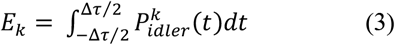

The resulting pulse energy, *E*_*k*_, represents the signal measured for a single pixel *k* in a given microscope, where 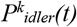is the time-dependent power of the idler pulse. Note that the output idler pulse is slightly shorter than the pump pulse (~1.7 vs. 2 ns) as FWM gain is nonlinear with respect to pump power (Eq. 2), leading to poorer amplification at the leading and trailing edges of the pump pulse. We then collect 100 of these single pixel measurements with a given signal power, *P*_*signal*_, to inform statistics for the amplification process (Fig. 3b) including the mean (*μ*_*signal*_) and standard deviation (*σ*_*signal*_) of the distribution. Comparing the statistics for a given signal power to the case where the signal is off (*μ*_*dark*_, *σ*_*dark*_), we can determine the SNR:

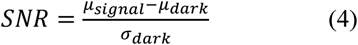

Note that *μ*_*dark*_ ≠ 0, as the amplification process inevitably creates ASE within the detection band. This ASE is the primary source of noise for the FOPA at high gain.

We next collect SNR measurements over a wide range of signal power (Fig. 3c). Assuming that the detector is primarily shot-noise limited, the SNR should be proportional to the square root of the amplified signal. Accordingly, we fit the SNR data on a logarithmic scale for all valid data points (*i.e*. SNR ≥ 1) to a line. The intercept of the line with an SNR of 1 corresponds to the signal power for which there is equal noise and signal detected, or the noise equivalent power (NEP). The NEP is effectively the threshold for detection, as noise and signal cannot be distinguished at powers below this value.

We next characterize the NEP as a function of amplifier gain, *G*, and bandwidth (Fig. 3d), where bandwidth is given by *Δv* = (2*Δτ*)^*–*1^. For low gain (*G* ~ 30 dB), the oscilloscope noise floor – or the readout noise – is the primary noise source and NEP is correspondingly large. As gain increases, the signal surpasses the noise floor, and NEP drops. For very high gain, there appears to be a slight increase in NEP in some of the measurements; this is likely due to ASE degrading the amplifier performance. The dynamic range of the oscilloscope limits the gain to < 50 dB, beyond which the idler pulses saturate the scope.

Considering Figure 3d, the NEP of the amplifier increases monotonically with increasing bandwidth. This is in keeping with the fact that NEP for a detector is typically specified by a single sensitivity value with units of W/Hz^1/2^. Accordingly, in Fig. 4, we plot minimum NEP as a function of bandwidth and fit the data to the function NEP = *NΔv* ^*1/2*^, where *N* is the detector sensitivity. We find that the sensitivities of the 50-m and 480-m amplifiers are 300 ± 90 fW/Hz^1/2^ and 200 ± 30 fW/Hz^1/2^, respectively, where the error bars are determined by the 95% confidence interval of each fit. For reference, we calculated the theoretical performance of a commercial NIR PMT (Hamamatsu H10330C-75, dashed pink line). The FOPA has comparable performance to the theoretical PMT sensitivity up to its bandwidth limit at ~100 MHz. We also included the shot-noise limit, the fundamental detection limit corresponding to the measurement of a single photon (black dashed line), given by *P*_*sn*_ = *hc/(λ*_*i*_*Δτ)* where *h* is Planck’s constant and *c* is the speed of light in vacuum. At the highest bandwidth tested (1.6 GHz), the NEP (~8.2 nW) is ~20-fold higher than the shot-noise limit (~0.4 nW), indicating that the FOPA has a sensitivity of ~20 photons/pixel at this speed.

**Fig. 4.**
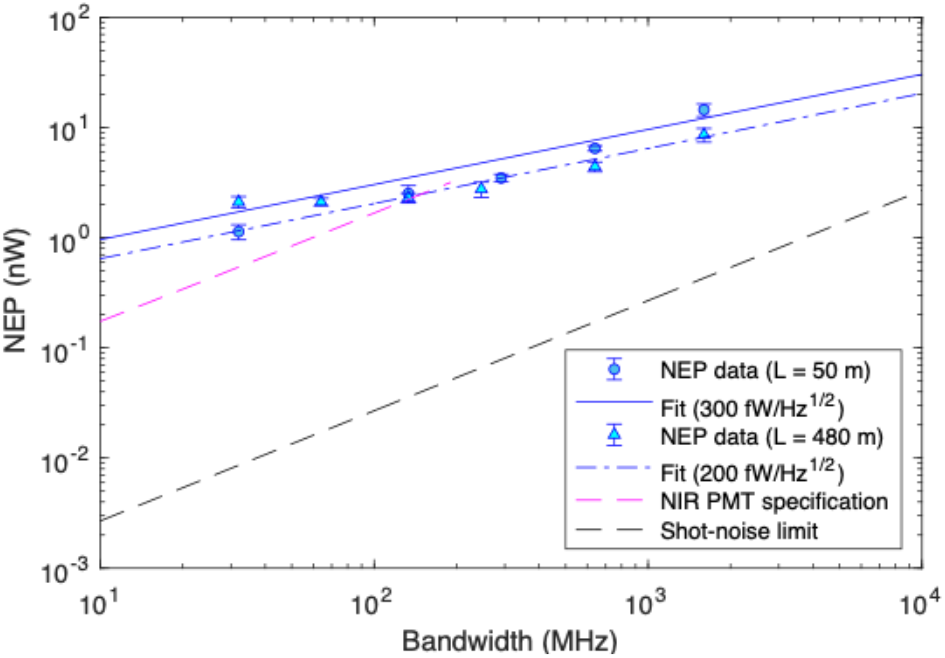
(a) NEP as a function of bandwidth for the 50-m (circles) and 480-m (triangles) amplifiers with fitting lines (solid and dot-dashed blue lines) corresponding to amplifier sensitivity; Theoretical sensitivity for a NIR PMT (Hamamatsu H10330C-75, pink dashed line) and the shot-noise limit (black dashed line).

To further benchmark FOPA performance, we conducted control experiments using the setup in Fig. 1c with two different TIAs (Femto DHCPA-100 using both the 80 MHz and 200 MHz bandwidth settings at high gain, Thorlabs TIA60 with 60 MHz bandwidth). Due to the finite bandwidth of the amplifiers, not all bandwidths tested for the FOPA (Fig. 4) could be replicated. Table 1 shows the results of the control experiments along with the NEP for each amplifier (50 and 480 m) extrapolated at the bandwidths for the control measurements using the amplifier sensitivities (300 fW/Hz^1/2^ and 200 fW/Hz^1/2^, respectively). The FOPA is ~100× more sensitive than the Femto amplifier, and ~10× more sensitive than the Thorlabs amplifier. These values are also borne out by the manufacturer specified sensitivity for each device: 155 and 5.8 pW/Hz^1/2^ in 200 and 80 MHz bandwidth modes, respectively, and 4.8 pW/Hz^1/2^ for the Thorlabs amplifier.

**Table 1.** NEP measurements using electrical amplification.

| Model | Bandwidth (MHz) | Electrical NEP (nW) | Optical NEP,<br>$L = 50 \text{ m}$ (nW) | Optical NEP,<br>$L = 480 \text{ m}$ (nW) |
| --- | --- | --- | --- | --- |
| Femto DHPCA-100,<br>80 MHz, high gain | 25 | 70 | 1 | 1.5 |
|  | 43 | 100 | 1.3 | 2 |
| Femto DHPCA-100,<br>200 MHz, high gain | 27 | 250 | 1 | 1.5 |
|  | 47 | 440 | 1.4 | 2.1 |
|  | 74 | 720 | 1.7 | 2.6 |
| Thorlabs TIA60 | 27 | 10 | 1 | 1.5 |
|  | 36 | 17 | 1.2 | 1.8 |

## 4. Confocal imaging with optical signal amplification

To test the capabilities of the FOPA for imaging experiments, we constructed the confocal microscope shown in Fig. 5. An ECL tuned to *λ*_*s*_ = 1620 nm served as the laser source and was coupled to the microscope through a 50:50 fiber beam splitter (the other port was optically terminated to avoid back reflections). The light was collimated onto a pair of galvanometric mirrors (Thorlabs GVS002) and relayed to the back focal plane of the objective lens (Thorlabs AC254-40-C) before being focused onto the sample. Back reflected light followed the same path in reverse and was de-scanned by the galvo mirrors before re-coupling to the fiber beam splitter. Only light that was reflected along the same trajectory from which it originated could mode match to the single-mode fiber and produce signal, which ensured the confocality of back reflected signals. Light sent back toward the ECL was blocked with an optical isolator, while the other port was connected to the signal input of the FOPA (*L* = 480 m, Fig. 1b), or, for control experiments, was directly detected by the PD and electrically amplified (Fig. 1c). Electrical signals from the control and FOPA conditions were fed into an FPGA (MBF Bioscience vDAQ) which also controlled the analog signals to run each galvo mirror.

**Fig. 5.**
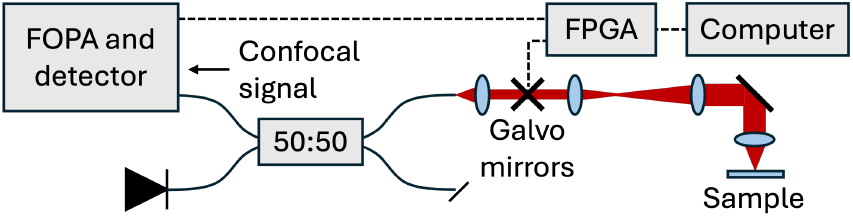
Confocal reflectance microscope; signal (*λ* = 1620 nm) is transmitted through a 50:50 fiber beam splitter to the microscope; scanning is facilitated by galvanometric mirrors; back-reflected signal from the sample goes back through the microscope, is de-scanned by the galvo mirrors, and re-coupled to the fiber splitter; signal transmitted backward to the upper port is coupled to the fiber optical parametric amplifier (FOPA) and detector (or directly to the detector and TIA for control experiments); amplified electrical signals are digitized by a field-programmable gate array (FPGA).

The bandwidth of the FPGA (<125 MHz) was not sufficient to capture the highest bandwidth capabilities of the FOPA. We accordingly elected to operate at 32 MHz (~16 ns pump pulses) for imaging experiments to ensure the FPGA bandwidth had no deleterious effects on the imaging results. Additionally, we limited control experiments to only the Thorlabs TIA60 amplifier, as it performed best in the previous control studies (Table 1). To stay within the safe, inertia-limited speed of the galvo mirrors, we slowed the frame rate down to ~0.3 Hz for each 1 Mpix image by increasing the pixel dwell time to ~3.3 μs (corresponding to ~13 pulses/pixel at a repetition rate of 4 MHz). In an optimized laser scanning system, the full FOPA bandwidth could be utilized with commercially available higher-bandwidth digitization hardware. Nonetheless, the experiments are a fair comparison between optical and electrical amplification.

To characterize the image quality provided by the confocal microscope with both electrical and optical amplifiers, we started by imaging a bright, well-defined sample consisting of a silver mirror with scratches in the coating. Figures 6a and 6b show the mirror under ~2 μW of signal power with optical and electrical amplification, respectively. For the same input power, the optically amplified image shows improved SNR and contrast. To quantify image SNR, we calculated the deviation of each pixel value, *I*_*k*_, from the image mean value (*μ*_*I*_) and normalized to the standard deviation of all pixels (*σ*_*I*_) before taking an average across all pixels, *N*_*p*_:

**Fig. 6.**
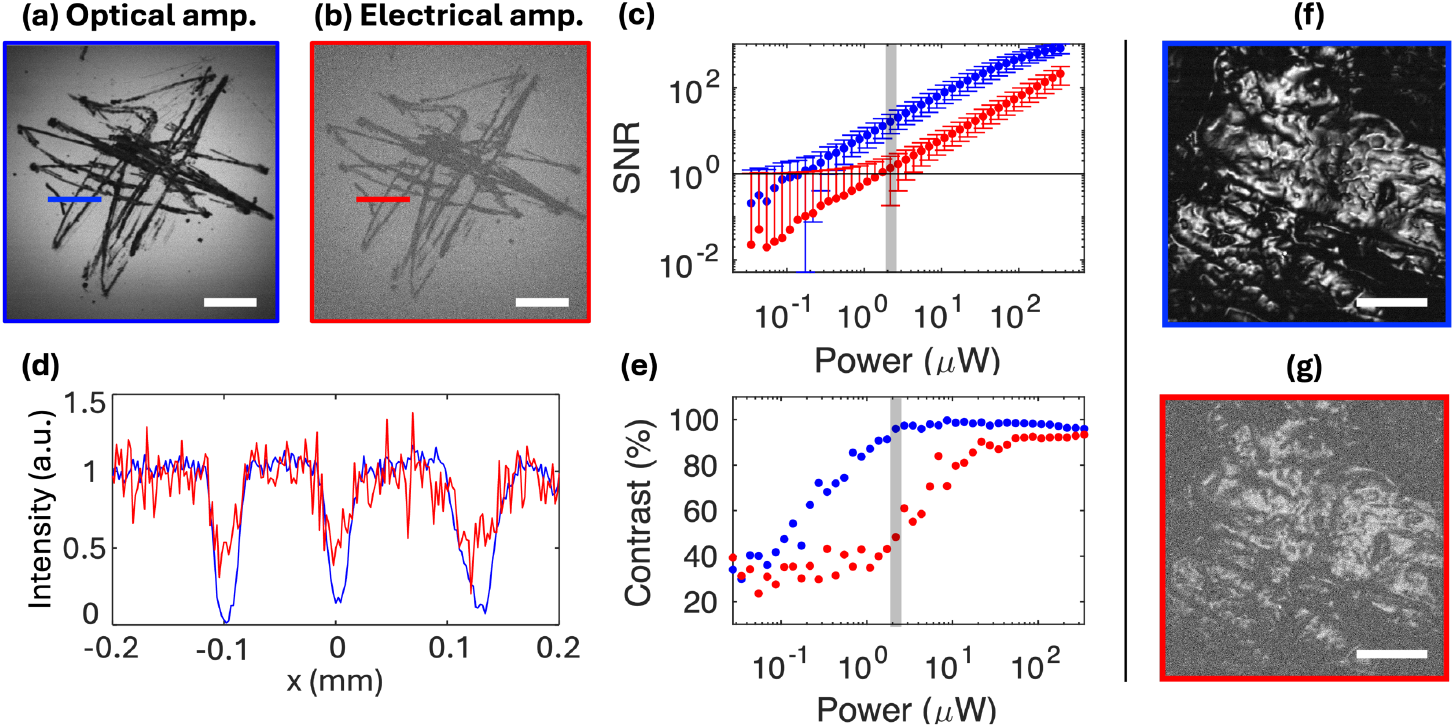
Images of a scratched silver mirror with optical (a) and electrical (b) amplification for 2.0 μW of signal power incident on sample; scale bar = 400 μm. (c) Mean image SNR as a function of signal power for electrical (red) and optical (blue) amplification; error bars correspond to standard deviation across pixels. (d) Line cuts of corresponding regions in (a) and (b) highlighting image contrast. (e) Calculated Michelsen contrast for line cuts per (d) as a function of signal power; in both (c) and (e) >10× more signal power is required to match optical performance with electrical amplification. Images of chicken muscle tissue using optical (f) and electrical (g) amplification for 55 μW of signal power incident on the sample; scale bar = 75 μm.

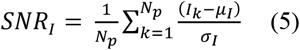

Figure 6c shows the image SNR as a function of signal power. At 2 μW (grey shading in Fig. 6c, highlighting the SNR values corresponding to the images in 6a and 6b), the SNR improves from 1.4 to 16.2 with optical vs. electrical amplification. This improvement factor of ~10× is consistent across the measurement. Additionally, we considered the contrast afforded by each amplification method by comparing features shared in each image (blue and red lines in Fig. 6a and 6b). Figure 6d shows the linecuts of these features, normalized to the same value for the high reflectivity portions of the line. The contrast clearly decreases for electrical amplification relative to optical amplification. We quantified contrast using the Michelsen definition: *C = (I*_*max*_*–I*_*min*_*)/(I*_*max*_*+I*_*min*_) [28], finding significant improvement in contrast for optical as opposed to electrical amplification (96% vs 48% for the 2 μW condition shown in Figs. 6a, 6b, and 6d). Looking closely, the trend in the contrast curve is roughly the same for both methods, however, optical amplification requires ~20× less power to achieve the same contrast as the electrical case.

Finally, we used the system to image chicken breast muscle tissue (*pectoralis major*). At the surface of the sample (Figs. 6f and 6g), the SNR and contrast are clearly improved in the optically amplified images relative to the electrically amplified case, in keeping with the rest of the characterization data. While the sample had no contrast agent that would facilitate imaging deeper in the tissue, we expect that the difference in performance between the two amplification methods would translate to increased penetration depth using the FOPA.

## 5. Discussion

The specifications (*i.e*. bandwidth, sensitivity, and dynamic range) of the point-detector is a bellwether for the functionality of a laser scanning microscope. Maintaining these crucial characteristics while operating at NIR wavelengths for compatibility with imaging deeper in turbid media has become a recent challenge in LSM, limited by the boundaries of conventional detector technology. NIR PDs are ideal detectors in terms of speed, operation wavelength range, and dynamic range, but are not responsive enough to produce signals above the detection noise floor of conventional digitization hardware for low-photon-flux applications.

Here, we demonstrate two optical amplifiers that bridge the responsivity gap between PDs and the small signals typical of LSM. The two different FOPA lengths (50 and 480 m) inform tradeoffs between pump power requirements and gain bandwidth, each providing >50 dB of parametric gain. The sensitivities for the amplifiers are ~300 and ~200 fW/Hz^1/2^, respectively, matching the theoretical performance of a NIR PMT up to its bandwidth limit of ~100 MHz. Both amplifiers outperform *post facto* electrical amplification of signals directly detected by a PD with ~10-100-fold increase in detection threshold. The maximum FOPA bandwidth (~1.6 GHz) exceeds that of both typical electrical amplifiers and NIR PMTs by a factor of ~10, with a NEP of ~8 nW, corresponding to a detection threshold of ~20 photons/pixel. In imaging experiments, we found that the improved performance of the FOPA was maintained, with optical amplification requiring >10× less power to achieve the same SNR and contrast as images obtained with electrical amplification.

While the optical detection properties outperform other conventional modalities, the detection threshold is ~10-fold higher than the shot-noise limit (Fig. 4). However, theoretically, FOPAs can detect signals as low as ~2 photons/pixel [24]. We believe that two areas of improvement could reduce the apparent excess noise. First, the loss and passband width of the filter used in the experiments could be improved by using a higher diffraction efficiency grating, or an etalon-based filter. The latter are available in all-fiber formats to remove free space elements from the system. Second, and crucially, the pump used for experiments has both high relative intensity noise (RIN) and ASE from the EDFA. Because the FOPA gain is nonlinear with respect to pump power (Eq. 2), the pump RIN is transferred to and amplified in the output idler and signal beams. Furthermore, power outside the pump wavelength due to ASE from the EDFA acts as a seed in the FOPA, which amplifies and increases output noise. However, we believe both issues would be greatly ameliorated by utilizing a CW, polarization-maintaining (PM) fiber laser to the pump the FOPA, rather than the non-PM system used for these experiments. We have seen in previous experiments that PM systems exhibit significantly less RIN than non-PM lasers, and CW rather than pulsed operation makes it easier to drive the amplifier at saturation and improve the signal extraction to reduce ASE in the pump beam.

In parallel, the apparent imaging quality improvements suggested by these imaging experiments can be benchmarked in LSM applications requiring detection of NIR signals. Pump-probe techniques such as MIP, SWIP, and CARS, among others, would be appropriate test grounds for this technique. Additionally, these experiments did not make use of the FOPA gain bandwidth (>30 nm full-width at half-maximum) – a trait that would be relevant for imaging NIR-fluorescing labels, or OCT. The results of these preliminary characterization and imaging experiments indicate that optical pre-amplification of low-photon-flux NIR signals is a viable approach to increase sensitivity, bandwidth, and dynamic range and represents a new paradigm for LSM with the potential to increase SNR, temporal resolution, and depth penetration for a wide variety of biomedical imaging applications.

## Data availability

Data underlying the results presented this paper are not available publicly at this time but may be obtained from the authors upon reasonable request.

